# A systematic screening assay identifies efficient small guide RNAs for CRISPR activation

**DOI:** 10.1101/2023.11.10.566523

**Authors:** Elin Arvidsson, Diana Duarte Lobo, Ermelinda Sabarese, Fabio Duarte, Rui Jorge Nobre, Cecilia Lundberg, Luis Quintino

## Abstract

CRISPR-mediated gene activation (CRISPRa) encompasses a growing field of biotechnological approaches with exciting implications for gene therapy. However, there is a lack of experimental validation tools for selecting efficient sgRNAs for downstream applications. Here, we present a screening assay capable of identifying efficient single- and double sgRNAs through fluorescence quantification in vitro. In addition, we provide a tailored Golden Gate cloning workflow for streamlined incorporation of selected sgRNA candidates into lentiviral (LVs) or adeno-associated vectors (AAVs). The overall workflow was validated using therapeutically relevant genes for neurodegenerative diseases, such as *Tfeb*, *Adam17*, and *Sirt1*. The most efficient sgRNAs also demonstrated activation of endogenous gene expression at mRNA and protein levels. Further proof-of-principle assays using *Tfeb* indicated that gene activation was accompanied by increased levels of Lc3b. This data demonstrates the potential of the screening assay to identify functionally efficient sgRNA candidates across multiple genes along with streamlined cloning of viral vectors and may assist in accelerating future developments of CRISPRa-focused applications.

## 1 Introduction

The discovery of clustered regularly interspaced short palindromic repeats (CRISPR) DNA segments and CRISPR-associated proteins (Cas), known as CRISPR/Cas, has revolutionized biomedical research (Katti et al., 2022). There is an ever-growing number of CRISPR/Cas systems engineered for genome editing and, interestingly, most of the CRISPR/Cas systems adapted for biotechnological purposes operate similarly. A Cas protein containing a small guide RNA (sgRNA) targets a nucleic acid sequence complementary to the binding sequence of the sgRNA and adjacent to a protospacer adjacent motif (PAM). The Cas/sgRNA complex then performs a double stranded break of the targeted region.

This modularity enables the adaptation of CRISPR/Cas systems to mediate programable genetic functions beyond gene editing. For example, CRISPR/Cas systems can perform specific epigenetic modulation, gene inhibition and gene activation (CRISPRa) (Kampmann, 2020). To adapt CRISPR/Cas systems for CRISPRa, the Cas catalytic domains are silenced and effector domains from transcription factors are fused to the silenced/dead Cas9 protein or bound to sgRNA via RNA loops. The Cas/sgRNA/effector complex targets a genomic DNA sequence and activates gene expression when bound upstream of transcriptional start sites (Gilbert et al., 2014).

Diverse CRISPRa systems have been previously described in the literature. Among them, CRISPRa-VPR (Chavez et al., 2015) has been widely used, and due to its reproducible gene activation now serves as a benchmark for testing new CRISPRa systems (Chavez et al., 2016; Zhang et al., 2018). The VPR effector domain consists of a fusion protein of VP64, p65 and Rta transcriptional effector domains. In a previous collaboration with the Church lab, our lab aided in the validation of a smaller version of the VPR domain (Vora et al., 2018) that together with a compact Cas could be used with viral vectors for gene therapy applications in vivo. The compact CRISPRa system was further engineered (MiniCas9V2) for applications in neurons and demonstrated activation of therapeutically relevant genes in the brain (Maria et al., 2020).

As gene therapies typically require the use of viral vectors, specifically lentiviral vectors (LV) and adeno associated vectors (AAV), the CRISPRa components need to be sufficiently small to be packaged into an LV or AAV. Accordingly, several factors need to be considered. First, a miniature Cas protein (Vora et al., 2018; Xu et al., 2021) is optimal. Second, a compact effector domain with proven capability of gene activation in vivo, such as SAM or VPR (Liu et al., 2021), should be tested. Lastly, the lowest possible number of sgRNAs should be used.

Presently, there are several apt Cas proteins and compact effector domains available for viral vector usage in vivo. However, there is a lack of in vitro screening methods for selection of functional sgRNAs to minimize the number of sgRNA needed for in vivo CRISPRa. In addition to reducing the size of the genetic payload that needs to be packaged into viral vectors, decreasing the number of necessary sgRNAs also minimizes the possible off target effects (Casas-Mollano et al., 2020; Schoger et al., 2020). Validating and selecting a minimal number of functional sgRNAs will further support the design of novel and sophisticated CRISPRa strategies. Without functional screening of sgRNA, up to four sgRNAs may be needed to ensure activation of target genes in vivo (Maria et al., 2020).

To optimize sgRNA for CRISPRa in vivo, it is important to have a fast, simple, and systematic experimental screening assay. Ideally, such an assay should be incorporated in a streamlined cloning workflow to generate LV and AAV quickly and efficiently. With these features in mind, a screening assay was designed using plasmid transfection and fluorescent protein readouts in vitro, coupling the plasmids used to a tailored Golden Gate cloning workflow to facilitate incorporation into LV and AAV. The screening assay was validated using therapeutically relevant genes for neurodegenerative diseases such as Alzheimer’s Disease, Parkinson’s Disease and Machado-Joseph Disease. Specifically, a disintegrin and metalloprotease 17 (*Adam17*) (Mockett et al., 2017), transcription factor EB (*Tfeb*) (Rai et al., 2021) and Sirtuin (*Sirt1*) (Matos et al., 2018) were targeted for activation.

The CRISPRa screening identified sgRNAs that led to robust activation of *Adam17*, *Tfeb* and *Sirt1*. To further validate the top sgRNAs in relevant cell lines, the sgRNAs were subcloned and transfected or transduced using LV. Most of the sgRNAs identified in the screening were able to activate endogenous gene expression, resulting in increased expression. It was further observed that gene activation was sufficient to elicit functional effects using *Tfeb* as an example. Taken together, these results suggest that the CRISPRa screening assay can identify single or dual sgRNA capable of strong activation of multiple therapeutically relevant genes. The described screening assay, together with the cloning workflow, should accelerate the development of novel CRISPRa applications for biomedical purposes.

## 2 Materials and Methods

### In silico small guide RNA design

To generate sgRNAs, 22-nt spacer sequences designed to target a 500-600 bp region immediately upstream of the *Tfeb*, *Adam17*, and *Sirt1* transcription start site (TSS). Spacer sequences were identified using the University of California Santa Cruz (UCSC) genome browser (www.genome-euro.ucsc.edu) and validated for template-strand targeting using the Benchling online setup tool (Benchling, 2023) (Benchling, n.d.)(Sup. Fig. 1). Seven binding sites were selected for each promoter, as this allowed single and dual sgRNA screening in a 96-well format. A 22-nt scrambled spacer (Scr) that did not target any sequence in the mouse genome, verified via BLAST, was generated as a control sgRNA used in all subsequent experiments. A standard S. aureus compatible RNA scaffold with a terminator sequence (gttttagtactctggaaacagaatctactaaaacaaggcaaaatgccgtgtttatctcgtcaacttgttggcgagattttttt) was added downstream of the sgRNA spacer sequence. The sequences used to generate nonpalindromic overhangs for assembly were previously described by the Barrick laboratory (Barrick, 2021). All visualization of DNA segments and assembly of constructs was performed in silico using SnapGene (Dotmatics).

### TOPO cloning

Constructs containing individual sgRNA segments, scrambled filler segments or destination blocks were designed as GeneArt™ Strings™ DNA fragments (ThermoFisher Scientific). The gene strings consisted of an expression cassette, dubbed Duplo (DPL0), containing an hU6-promoter, sgRNA binding site, *S. aureus* sgRNA scaffold and poly-T terminator sequence. The DPL0 cassette was flanked by a 4-bp overhang sequence and *Esp3I* recognition sites to generate non-palindromic nucleotide overhangs upon restriction. Filler blocks were designed as random 400 nt segments similarly flanked by a 4-bp overhang sequence along with *Esp3I* restriction sites to ensure that the downstream Golden Gate cassettes were similar in size. In addition, a Lego (LG0) cassette consisting of multiple Golden Gate destination sites and standard multiple cloning sites was also designed for downstream assembly. Gene strings were inserted into a pCR®II-TOPO® vector using the Zero Blunt™ TOPO™ PCR Cloning Kit according to the manufacturer’s instructions (Invitrogen) to generate pDPL0, pScramble and pLG0 constructs, respectively.

### Gateway Cloning

Gateway 4.1.2 assembly was used to generate pHG.EF1a.MiniCas9V2, pHG.mTfeb.TdTomato, pHG.mAdam17.TdTomato and pBG.mSirt1.TdTomato reporter constructs. All plasmids containing the promoter regions were ordered via gene synthesis as Gateway-compatible donor plasmids (ThermoFisher Scientific). Promoter sequence and expression cassette from donor constructs pEntry.P4P1R.EF1a and pEntry.MiniCas9V2 (Maria et al., 2020) were assembled in the pHG destination vector (Quintino et al., 2013) to generate the pHG.EF1a.MiniCas9V2 construct. For the TdTomato reporters, donor vectors pEntry.P4P1.mTfeb, pEntry.P4P1R.mAdam17 or pEntry.P4P1.mSirt1, along with pEntry.TdTomato, were assembled in pHG or pBG destination lentiviral transfer vectors. Reactions were performed using the Gateway™ LR Clonase™ II Enzyme mix (Invitrogen) according to manufacturer’s directions.

### Golden Gate Cloning

Expression cassettes were excised from the pDPL0 vector through *Esp3I-*digestion and ligated in tandem using T4 ligase (20U/µl, New England Biolabs) for assembly into the *Esp3I*-digested pLG0 vector backbone. Inserts and backbone were added to the reaction mix at a 1:1 insert:vector ratio using 20 fmol of each plasmid and supplemented with 50mM ATP and 100mM DTT. Assembly was performed at a 20 µl final reaction volume. Thermal cycling was programmed for 25 cycles of digestion at 37°C for 2 min and ligation at 16°C for 5 min, followed by *Esp3I* digestion at 60°C for 10 min and finalization by heat inactivation at 80°C for 20 min. This cloning was performed in accordance with protocols previously established by Haynes and Barrett (Haynes and Barrett, 2013).

### Standard Molecular Cloning

Assembly cloning was used to insert expression cassettes into AAV- or lentiviral compatible vectors. pLG0-constructs and the pAAV-U6-mAtct1-Sa acceptor vector (Vora et al., 2018) were digested with *NotI-MfeI*, while the pHG.EF1a.mRFP acceptor vector was digested with *SpeI*. Digested products were separated on a 1% agarose gel. Bands corresponding to the expression cassette and digested acceptor vector, respectively, were excised and purified using the GeneJET Gel Extraction Kit (ThermoFisher Scientific) or the NucleoSpin Gel and PCR Clean-up Kit (Macherey-Nagel) according to the manufacturer’s instructions. Purified DNA concentrations were quantified against a GeneRuler 1kb Plus DNA ladder (ThermoFisher Scientific) and ligation was performed at a 3:1 insert:vector ratio using Anza^TM^ T4 DNA Ligase Master Mix (ThermoFisher Scientific) supplemented with 10mM ATP. The ligation reaction was performed according to manufacturer’s instructions.

### Sequencing

Finished constructs were Sanger sequenced through the Eurofins TubeSeq service (Eurofins Genomics). M13-forward and reverse universal primers were used to sequence the pDPL0 and pLG0 constructs.

### Transformation

Bacterial transformation was performed in One Shot^TM^ TOP10 (Invitrogen) for pDPL0 and pLG0 or Stbl3^TM^ competent bacteria (Invitrogen) for the LV and AAV plasmid backbones, respectively. Reactions were performed according to manufacturer’s instructions, using 1 µl of ligated plasmid per reaction. One hundred microlitres of transformation mix was plated on antibiotic-supplemented LB agar plates and incubated overnight at 32°C or 37°C, respectively.

### DNA extraction

Colonies from culture plates were selected and grown in 5 ml antibiotic-supplemented LB medium overnight (16h). DNA was extracted using the QIAprep Spin Miniprep Kit (QIAGEN, Germany). Control restriction was performed using standard methods and correct clones were selected and grown overnight in 250 mL antibiotic-supplemented LB medium. The correct bacterial cultures were processed the following day using the NucleoBond Xtra Midi kit for transfection-grade plasmid DNA (Macherey-Nagel) according to the manufacturer’s instructions. Purified DNA was resuspended in molecular grade water to prevent downstream cloning inefficiency due to the presence of EDTA in standard resuspension buffer, especially when performing Golden Gate reactions.

### Cell culturing and transfection for CRISPRa screening assay

Tissue culture plates were incubated with sterile water containing 0.002% Poly-L-Ornithine Solution (PLO) (Merck) overnight, washed twice with sterile water or PBS and left to dry fully in a ventilated hood before cell seeding. 293T cells, maintained in Dulbecco’s Modified Eagle Medium (DMEM) (Invitrogen), supplemented with 10% fetal bovine serum (FBS) and 1% Penicillin-Streptomycin (P/S) (Invitrogen), were seeded at a density of 6000 cells per well. For transfection, pHG.EF1a.BFP, pHG.EF1a.TdTomato carrying the promoter of the gene of interest, pHG.EF1a.MiniCas9V2 and pDPL0 sgRNA constructs were resuspended in PBS at a 1:0.1:2:1 ratio to a total DNA content of 200ng/well. Polyethyleneimine 25,000 (PEI) (1 mg/ml) was added to the plasmid mixture at a 5:1 PEI:DNA ratio and incubated for 15 min at RT before addition to the cell culture. Transfected cultures were incubated at 37°C for 48h before media was aspirated. Cells were fixed with 4% PFA for 10-15 min at RT before being washed twice with PBS. The fixed cultures were quantified as described in the fluorescent analysis section below.

### Gene expression assay

Neuro2A cells were maintained in culture with DMEM (Sigma-Aldrich) supplemented with 10% FBS and 1% P/S, or with 50% DMEM high-glucose (Cytiva) + 50% Opti-MEM (Gibco) supplemented with 5% FBS and 1% P/S. Cells were seeded at a density of 300,000 cells/well in 6-well culture plates. Twenty-four hours later, transfection was performed using PEI MAX 40,000 (1 mg/ml) at 7.5:1 PEI:DNA ratio. pHG.EF1a.MiniCas9V2 was added along with the respective sgRNA candidates at a 2:1 ratio to a total of 600 ng DNA per well. The transfection mixture was vortexed and incubated for 15 min at RT before being added to the cell culture.

Fourty-eight hours later, cells were harvested and processed using the NucleoSpin RNA kit (Macherey-Nagel) or the PureLink RNA Mini Kit (ThermoFisher Scientific) according to manufacturer’s instructions. cDNA was synthesized from 1 µg of RNA per sample using the iScript cDNA Synthesis Kit (Bio-Rad) or iScript Reverse Transcription Supermix for RT-qPCR (Bio-Rad) according to manufacturer’s instructions. For *Tfeb* and *Adam17*, reactions were set up at 10 µl reaction volume using LightCycler 480 SYBR Green Master Mix (Roche) and performed using the LightCycler 480 Real-Time PCR Instrument (Roche). The amplification protocol was performed as follows: pre-incubation at 95°C for 5 min, followed by 45 cycles of amplification including denaturation at 95°C for 10s, followed by annealing at 60°C for 10s, and extension at 72°C for 10s. The melting curve was performed at 65°C and increased at a ramp rate of 4.8°C/seconds up to 95°C with a hold time of 5 minutes. For *Sirt1*, quantitative real-time PCR was carried out at 20µl reaction volume and performed in CFX96 Touch Real-Time PCR Detection System (Bio-Rad). The amplification protocol started with denaturation step at 95°C for 30s, followed by 40 cycles of two steps: denaturation at 95°C for 5s and extension/annealing at 60°C for 15s. The melting curve was performed at 65°C for 5s, and up to 95°C with an increment of 0.5°C. Results were analyzed in terms of mRNA quantification relative to control samples and determined by the Pfaffl method (Pfaffl, 2001). Values were analyzed using standard delta-delta-Ct calculations.

The following primers were used to amplify genomic targets of interest: *Tfeb* (forward: AGCAGGTGGTGAAGCAAGAG, reverse: TTGGACAGGTTGGGGAATGG), *Adam17* (forward: GTGGCTCTCAACTCTGTAACTC, reverse: TTTACAGCACTTGGCTTTGTTT), *Sirt1* (forward: AGCGGCTTGAGGGTAATCA, reverse: ACTGCCACAGGAACTAGAGGA), and SaCas9 (forward: CTGCTGAACAACCCCTTCAAC, reverse: TTGCTGATTCTGCCCTTGCC).

### Lentivirus production

Production of lentiviral constructs was adapted from previously described methods (Quintino et al., 2013). 12.5×10^6^ HEK 293T cells were seeded in 24 ml DMEM (10% FBS, 1% Pen/Strep) in a T_175_ surface-treated cell culture flask. On the following day, culture medium was replaced with 16.2 ml fresh culture medium. pHG.EF1a.(sgRNA).mRFP, pHG.EF1a.MiniCas9V2, pHG.EF1a.mRFP, or pHG.EF1a.Scramble.mRFP respectively were packaged using a p8.91 packaging plasmid, kindly gifted by Simon Davis (RRID:Addgene_187441) and pseudotyped using a pMD2.G VSV-G envelope expression plasmid, kindly gifted by Didier Trono (RRID:Addgene_12259).

Envelope, packaging, and transfer plasmids were resuspended in PBS at a 1:2:4 ratio, and PEI was added dropwise at a 3:1 PEI:DNA ratio. The transfection mixture was briefly vortexed and incubated at RT for 15 minutes before 1.8 ml transfection mixture was added gently to the medium of each culture flask. Two T_175_ flasks were prepared for each viral batch. After 48h, the medium of two T_175_ flasks per viral batch was pooled and centrifuged at 800xg for 10 min at 4°C. The supernatant was filtered through a 0.45 µm filter and centrifuged at 111,000xg for 90 min. Viral pellets were resuspended in 80 µl PBS and incubated overnight at 4°C before final resuspension, aliquoting and storage at −80°C. Titration of viral constructs was performed as described previously (Quintino et al., 2013).

### Lentiviral transduction

NIH-3T3 cells (ECACC nr. 93061524) were cultured at 6000 cells per well in DMEM (+10% FBS, +1% P/S) on Perkin Elmer LLC CellCarrier-96 Ultra Microplates (Perkin Elmer) coated with PLO overnight, as described above. Transductions were performed at an MOI of 15 immediately after seeding. All wells that did not contain sgRNA, aside from mock controls, were transduced with a pHG.EF1a.mRFP construct as a general cell marker to estimate transduction efficiency. Briefly, LV-expressing MiniCas9V2 and LV-expressing sgRNA together with mRFP were delivered to NIH-3T3 cultures at an MOI of 15. Seventy-two hours later, the samples were stained for Tfeb and the autophagosomal marker Lc3b. mRFP was used to identify transduced cells. Trehalose was used as a positive *Tfeb* activation control. Fourty-eight hours post-transduction, the media for selected cultures transduced with LV expressing mRFP was replaced with fresh media containing 100mM trehalose. Cells transduced with LV expressing mRFP alone, LV expressing Scr-sgRNA together with mRFP, and LV expressing MiniCas9V2 were used as additional controls.

### Immunofluorescence

Seventy-two hours after seeding NIH-3T3 cells, the media was aspirated and the cells were fixed in 200 µl of 2% PFA solution for 15 min and washed twice with PBS. Subsequently, cultures were prepared according to an in-house protocol for immunocytochemistry. Briefly, cultures were washed three times with 20mM KPBS (3.67 mM KH_2_PO_4_, 17 mM K_2_HPO_4_, 157 mM NaCl, pH 7.4) and incubated with 200 µl blocking solution (5% goat serum, 0,25% Triton-X in KPBS) at RT for 1h.

Primary antibodies, Rabbit anti-TFEB (ProteinTech) or Rabbit anti-LC3B (MilliporeSigma), were diluted 1:200 in 200 µl blocking solution per well, and added to the cells for overnight incubation at 4°C. The next morning, the primary solution was aspirated, and the cells were washed three times with KPBS, blocked for 1h at RT, and incubated with secondary antibody (Goat anti-Rabbit IgG (H+L) Cross-Adsorbed Secondary Antibody, Alexa Fluor 647 (Invitrogen)) and DAPI (Sigma Aldrich) diluted 1:1000 in 200 µl blocking buffer per well at RT for 1h. Cells were washed twice with KPBS and left in 200 µl KPBS supplemented with 0.1% NaN_3_ for preservation.

### Endogenous fluorescence and immunofluorescence analysis

For *Tfeb* and *Adam17*, activation was estimated though fluorescence quantification using a Plate RUNNER HD (TROPHOS). Alignment was performed using the Align software and fluorescent images of TagBFP2 (381/445 nm) and TdTomato (554/581 nm) expression were acquired using the Goelan software. Image analysis was performed using Image J (NIH). The total TdTomato fluorescence per plate was normalized by the total TagBFP2 fluorescence to adjust for transfection efficiencies. The mean TdTomato/TagBFP2 ratio of scramble samples was used as a baseline for subsequent calculations.

For *Sirt1*, Operetta CLS (PerkinElmer) was used for microplate fluorescence quantification. In brief, TagBFP2 cells were identified and TdTomato fluorescence in these cells was subsequently measured. The average sum per well of TdTomato fluorescence per cell was normalized by the average sum per well of TagBFP2 fluorescence per cell. The mean TdTomato/TagBFP2 ratio of Scr samples was used as baseline for subsequent calculations.

For *Tfeb* endogenous activation and Lc3b function assessment, cells were detected based on nuclear DAPI staining and a nuclear area of >50 µm^2^, <400 µm^2^. mRFP-positive cells were selected based on a mean nuclear mRFP intensity of >600 arbitrary units determined through scatterplot analysis.

Alexa-647 intensity within mRFP-positive cells was quantified to measure Tfeb and Lc3b expression and the mean of the pixel sums for Tfeb or Lc3b was measured. More extensive Operetta CLS parameters for *Tfeb* endogenous activation can be found in the supplementary tables.

### Synergy estimation

Effect-based methodology was used to assess synergy or antagonism between sgRNA. The significance of sgRNA interaction was first determined using two-way ANOVA (Slinker, 1998). For example, for each sgRNA combination, the activation of Scr (control), activation of single sgRNA A, activation of single sgRNA B and activation of the sgRNA A+B combination were evaluated together. The null hypothesis was that activation of sgRNA A was independent from activation with sgRNA B. Whenever there was a statistically significant interaction effect between activation of sgRNA A and sgRNA B, the null hypothesis was rejected. In other words, the activation was most likely a result of a significant interaction between sgRNA A and sgRNA B. To determine if the nature of the interaction between sgRNA A and sgRNA B was positive (synergy) or negative (antagonism), the two-way ANOVA interaction results were analysed together with a Combination Index (CI), which, due to the nature of the data, was calculated using a response additivity model (Foucquier and Guedj, 2015). The expected additive effect, Expected activation (A+B), was calculated by adding the activation of sgRNA A and activation of sgRNA B, as shown by the following equation: *E act* (*A* + *B*) = *act sgRNA A* + *act sgRNA B*. The CI was calculated by dividing the Expected activation of sgRNA A and sgRNA B by the observed activation of combining sgRNA A and sgRNA B, as shown by the following equation: 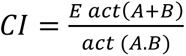. In synergistic sgRNA interactions CI < 1, in additive or independent interactions CI = 1 and in antagonistic interactions CI > 1. By determining if sgRNA A and sgRNA B had a significant interaction and using the CI to estimate if this interaction was positive or negative, it was possible to infer significant synergistic or antagonistic interactions between sgRNA combinations.

### Statistics

Statistical analysis was performed in GraphPad Prism 9 (Dotmatics). Experimental samples were compared against a scrambled control construct using multiple comparison ordinary One-Way ANOVA. Dunnett’s post hoc analysis was performed, using the Scr + MiniCas9V2 condition as control for comparison. Two-way ANOVA was used to calculate significant interaction scores. Histograms are presented as mean ± SEM.

## 3 Results

### L3G0 cloning workflow allows efficient and seamless assembly of multiple sgRNA combinations

The purpose of this study was to create an experimental screening system for sgRNA used for CRISPRa applications requiring viral vectors. Moreover, to make it truly universal in usage, the screening assay was coupled to a cloning workflow (Fig. 1). This enables a streamlined validation process, from plasmid design to functional assessment of endogenous gene activation. The initial task involved establishing a robust and flexible cloning methodology. This tailored Golden Gate workflow, named **L**und **E**fficient s**g**RNA **O**mnibus (L3G0) cloning, was designed in-house to provide a time- and labour-efficient strategy for subcloning multiple relevant sgRNA combinations into a single pLG0 plasmid that could then be transferred to plasmids containing LV or AAV backbones, according to specific research needs (Fig. 2).

**Figure 1:**
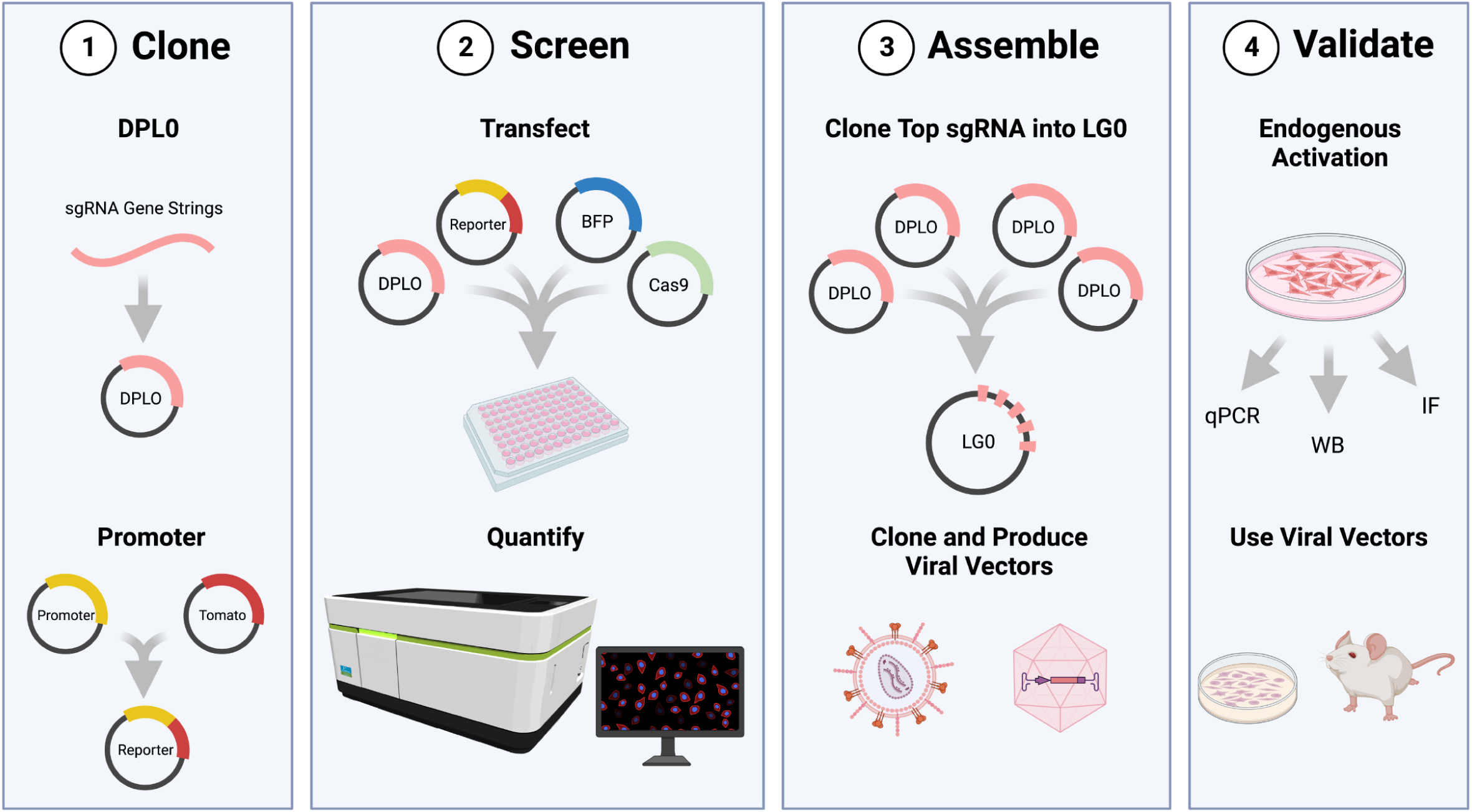
Workflow of the screening assay. 1) DPL0 cassettes containing candidate sgRNA sequences are cloned into TOPO vectors, as well as the reporter plasmid consisting of TdTomato driven by the promoter sequence for the gene of interest. 2) pDPL0 vectors are transfected in combination with MiniCas9V2, reporter, and BFP for normalization of fluorescence expression. Levels of TdTomato expression are normalized by BFP and quantified to identify transcriptionally efficient sgRNA combinations. 3) Selected cassettes are in turn assembled in pLG0 constructs and subcloned into LV or AAV, according to research needs. 4) Final validation of sgRNA and viral vectors and subsequent use in primary cells or in vivo. Figure was created in BioRender.com.

**Figure 2:**
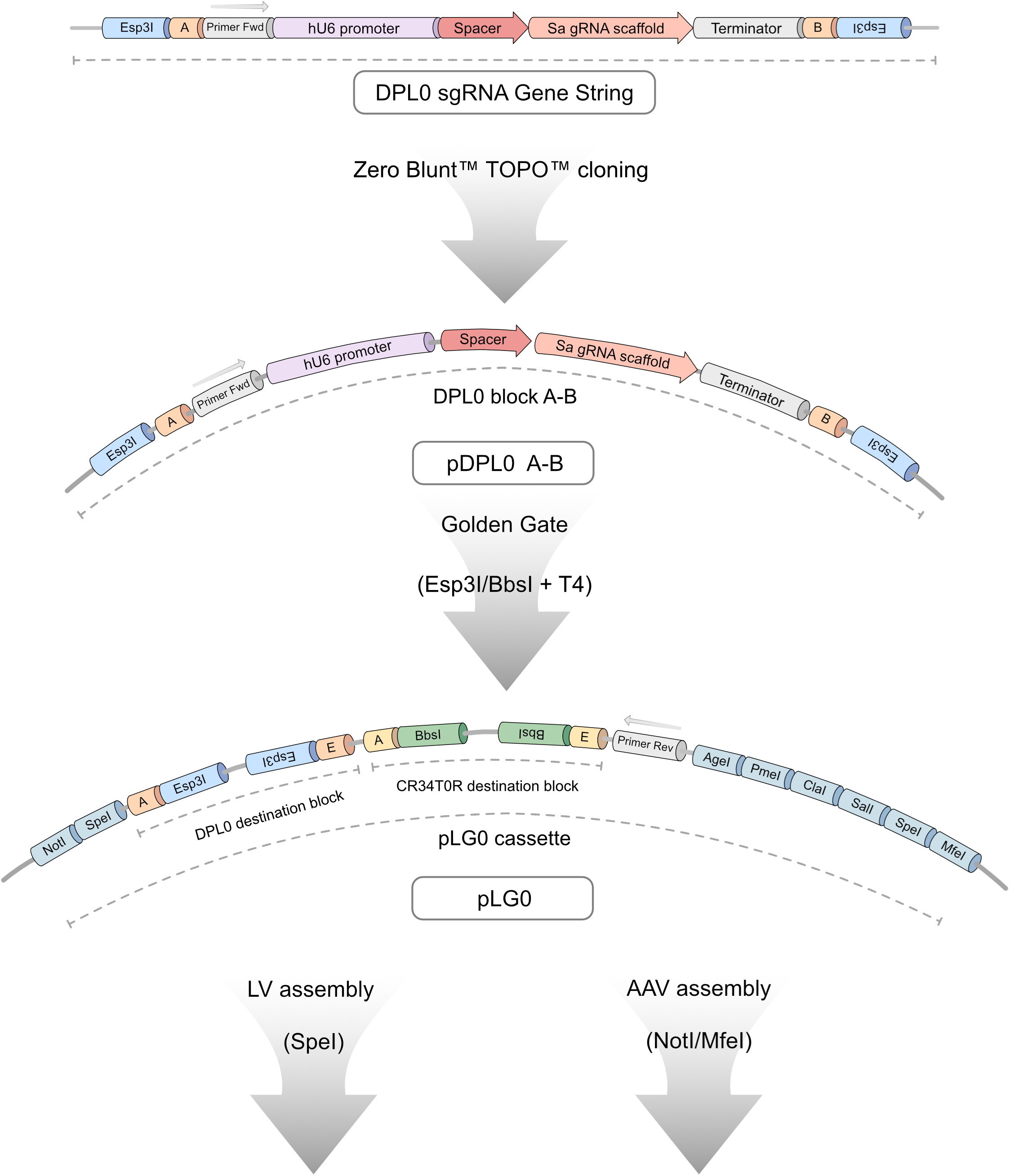
Detailed overview of L3G0 cloning workflow. All sgRNA candidates are ordered as a Gene String consisting of a DPL0 cassette containing a human U6 promoter, spacer or binding site, SaCas9 RNA scaffold, and downstream terminator sequence. The expression cassette is flanked by Esp3I-restriction target sites, generating 4-bp overhang sequences compatible with ligation of four separate DLP0 blocks in tandem through Golden Gate cloning. Ligated cassettes are introduced into destination blocks on the pLG0 construct that contain Esp3I-combatible sites (DPL0 destination) and BbsI-compatible sites (CR34T0R destination). This design allows seamless assembly of up to eight sgRNA candidates within a single vector. When in pLG0, primer binding sites allow for screening using colony PCR. The LG0 cassette provides a multitude of additional restriction sites, allowing for downstream applications such as subcloning into LV- and AAV-compatible constructs.

CRISPRa-compatible sgRNA binding sequences were selected using online bioinformatic tools such as Benchling and CHOPCHOP (Benchling, 2023) (Labun et al., 2019) to target the 500-600 base pair promoter region immediately upstream of the TSS. This region was selected as previous studies have highlighted this genomic window as suitable for CRISPRa sgRNA targeting (Gilbert et al., 2014; Konermann et al., 2015). As *Tfeb*, *Adam17* and *Sirt1* were selected for proof-of-concept validation, the mouse *Tfeb*, mouse *Adam17* and mouse *Sirt1* promoter regions were used. In total, seven sgRNA candidate binding sequences for each gene of interest were selected for insertion into DPL0 expression cassettes subsequent screening analysis (Sup. Fig. 1).

### CRISPRa screening assay determines experimental activation of promoters through fluorescence quantification

The CRISPRa screening assay was designed to experimentally estimate the activation efficiency of sgRNA identified using bioinformatic tools. The screening assay has four components: pDPL0 plasmids containing a sgRNA candidate, a plasmid expressing MiniCas9V2 (Maria et al., 2020), a reporter plasmid expressing TdTomato under the control of the promoter regions of interest, and a plasmid expressing TagBFP2 to be used as transfection control. The assay was performed by transfecting these four types of plasmids into 293T-cells and assessing fluorescence 48 hours after transfection. TagBFP2 expression was used as a transfection normalizer and the TdTomato expression in each condition was then compared to the mean ratio of a scramble (Scr) sgRNA control. Representative fluorescence images obtained using the *Sirt1* screening assay, can be found in Sup. Fig. 2.

One-way ANOVA demonstrated statistically significant differences in mouse *Tfeb* promoter activation (Fig. 3A) between sgRNA groups (F (28,58) =6.34, p<0.0001). Post hoc analysis with Dunnett test using the Scr condition as control showed that sgRNA 6 (3.45 ± 0.40 fold), sgRNA 1+3 (3.15 ± 0.23 fold), sgRNA 3+4 (3.96 ± 0.10 fold), sgRNA 4+5 (3.02 ± 0.83 fold), sgRNA 4+6 (3.40 ± 0.16 fold), sgRNA 5+6 (3.19 ± 1.09 fold) and sgRNA 5+7 (3.46 ± 1.02 fold) led to significant increases in TdTomato levels, indicating that these seven respective conditions resulted in robust mouse *Tfeb* promoter activation.

**Figure 3:**
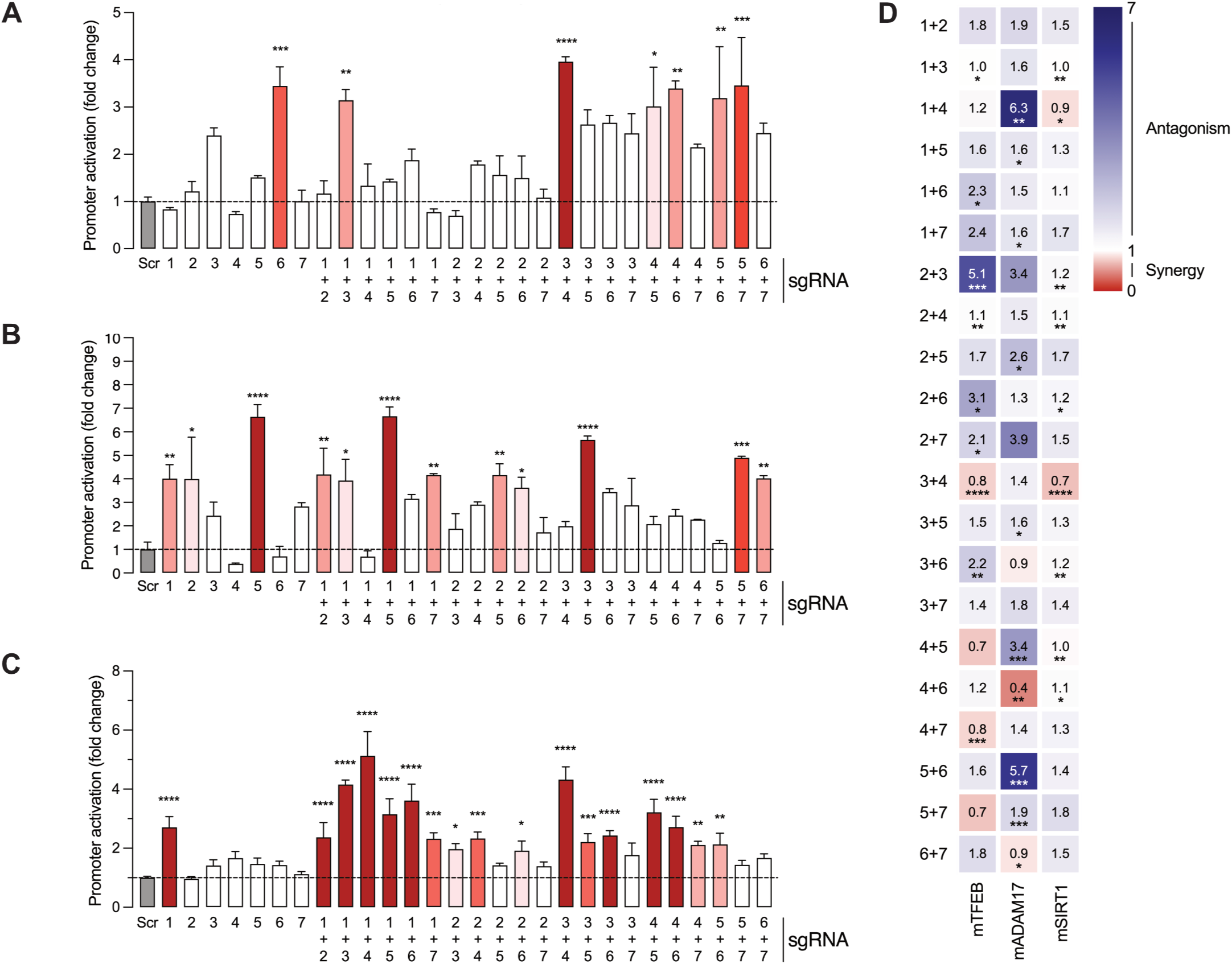
Screening assay identifies sgRNAs capable of activating transcription from *Tfeb*, *Adam17* and *Sirt1* promoters. 293T cells were transfected with pDPL0 expressing sgRNA, reporter constructs expressing TdTomato, MiniCas9V2 and BFP. Fourty-eight hours after transfection, the cells were fixed and the TdTomato and BFP fluorescence was measured. A) Activation of *Tfeb* expression by single or dual sgRNA combinations. B) Activation of *Adam17* expression by single or dual sgRNA combinations. C) Activation of *Sirt1* expression by single or dual sgRNA combinations. All values are presented as themean ± SEM and normalized against a Scr control sample (left, grey). For the histograms, bars with higher statistical significance are marked in darker shades of red. D) Calculation of synergy or antagonism for dual sgRNA combinations using two-way ANOVA interaction scores and Combination Index. Scr-Scramble. *P≤0.05, **P≤0.01, ***P≤0.001, ****P≤0.0001.

Similarly, one-way ANOVA showed statistically significant differences in mouse *Adam17* promoter activation (Fig. 3B) between groups (F (28,58) = 8.2, p<0.001). In the case of mouse *Adam17* sgRNA candidates, Dunnett post hoc test identified significantly increased promoter activation in a total of twelve combinations, including sgRNA1 (4.01 ± 0.59 fold), sgRNA 2 (4.00 ± 1.78 fold), sgRNA 5 (6.64 ± 0.52 fold), sgRNA 1+2 (4.19 ± 1.12 fold), sgRNA 1+3 (3.93 ± 0.91 fold), sgRNA 1+5 (6.66 ± 0.39 fold), sgRNA 1+7 (4.15 ± 0.06 fold), sgRNA 2+5 (4.16 ± 0.48 fold), sgRNA 2+6 (3.63 ± 0.45 fold), sgRNA 3+5 (5.66 ± 0.16 fold), sgRNA 5+7 (4.90 ± 0.06 fold), and sgRNA 6+7 (4.03 ± 0.11 fold).

In the case of mouse *Sirt1*, one-way ANOVA indicated statistically significant differences in mouse *Sirt1* promoter activation between groups (F (28,58) = 30.5, p<0.0001, Fig. 3C). Dunnett’s post hoc test indicated increased promoter activation in seventeen conditions, namely sgRNA1 (2.7 ± 0.21 fold), sgRNA 1+2 (2.37 ± 0.29 fold), sgRNA 1+3 (4.16 ± 0.09 fold), sgRNA 1+4 (5.13 ± 0.47 fold), sgRNA 1+5, (3.15 ± 0.30 fold), sgRNA 1+6 (3.62 ± 0.32 fold), sgRNA 1+7 (2.3 ± 0.11 fold), sgRNA 2+3 (1.97± 0.10 fold), sgRNA 2+4 (2.33 ± 0.12 fold), sgRNA 2+6 (1.91± 0.19 fold), sgRNA 3+4 (4.32± 25 fold), sgRNA 3+5 (2.20 ± 0.16 fold), sgRNA 3+6 (2.42 ± 0.10 fold), sgRNA 4+5 (3.20 ± 0.25 fold), sgRNA 4+6 (2.72 ± 0.21 fold), sgRNA 4+7 (2.10 ± 0.07 fold) and sgRNA 5+6 (2.13 ± 0.22 fold)

Although it was expected that not all single sgRNA or dual sgRNA screened would consistently lead to increased promoter activity, the number of single or dual sgRNA combinations capable of significant gene activation varied between the different promoter regions, ranging from seven for mouse *Tfeb* to seventeen for mouse *Sirt1*. Additionally, varying combinations of sgRNA led to notably higher levels of activation. These findings suggest that efficient sgRNA binding and gene activation are promoter specific.

To further investigate possible interactions between sgRNA in the target promoters, significant synergistic or antagonistic activation effects were calculated for the different sgRNA combinations in all the genes tested (Fig. 3D). In the case of *Tfeb*, combinations of sgRNA 1+6, sgRNA 2+3, sgRNA 2+4, sgRNA 2+6 and sgRNA 3+6 displayed an antagonistic interaction. In contrast, combinations of sgRNA 3+4 and sgRNA 4+7 had a synergistic interaction. For *Adam17*, sgRNA 1+4, sgRNA 1+5, sgRNA 1+7, sgRNA 2+5, sgRNA 3+5, sgRNA 4+5 and sgRNA 5+6 had an antagonistic interaction, while sgRNA 4+6 and sgRNA 6+7 exhibited a synergistic interaction. Finally, in the case of *Sirt1*, sgRNA 2+3, sgRNA 2+4, sgRNA 2+6, sgRNA 2+7, sgRNA 3+6, sgRNA 4+5, and sgRNA 4+6 had an antagonistic interaction, whereas sgRNA 1+4 and 3+4 had a synergistic interaction.

### Subcloning sgRNA into LV

After determining sgRNA candidates capable of robust activation, the selected multiple sgRNA cassettes present in the pDPL0 plasmids were assembled into the pLG0 plasmid (Fig. 2), as part of the L3G0 cloning workflow. This plasmid contains the LG0 cassette, consisting of two independent Golden Gate destination blocks surrounded by multiple cloning sites that were tailored to our LV- and AAV-compatible plasmids. The Golden Gate DPL0 destination block present in the pLG0 plasmid uses Esp3I for inserting sgRNA cassettes present in pDPL0 plasmids, whereas the CR34T0R destination block uses BbsI to clone in sgRNA-compatible cassettes. These two Golden Gate destination cassettes were designed for seamless tandem insertion of up to eight sgRNA cassettes within a single vector. More sgRNA cassettes can be added by designing more overhangs within the DPL0 and CR34T0R blocks (Potapov et al., 2018) or inserting additional pre-assembled DPL0 or CR34T0R groups of sgRNA cassettes in the downstream multiple cloning site region. Even with both destination blocks in use, the LG0 cassette reaches a size of 3.4kb, well below the packaging limit of an AAV. In the final part of the L3G0 cloning workflow, the multiple sgRNA cassettes present within the LG0 cassette are then subcloned into compatible LV or AAV.

### Selected sgRNA candidates lead to increased endogenous gene activation

After assembling selected sgRNA candidates into pLG0 plasmid backbones, the next step was to validate the ability of the single sgRNA or dual sgRNA combinations to activate endogenous gene expression. Selected pLG0 were transfected together with MiniCas9V2 in Neuro2A cell cultures. Transfection control consisted of cells transfected with PEI alone. After 48 hours, samples were harvested for RNA isolation. Samples were quantified by normalizing the expression levels of the respective gene of interest, using constitutively expressed dCas9 as transfection control and normalizer.

One-way ANOVA displayed significant differences between groups in samples transfected with *Tfeb* sgRNA in combination with MiniCas9V2 (F (3, 12) = 12.27), p<0.005). Dunnett’s post hoc analysis revealed that samples transfected with *Tfeb* sgRNA 6 displayed an over two-fold increase in *Tfeb* expression (2.10±0.25-fold) when compared to the Scr control (Fig. 4A).

**Figure 4:**
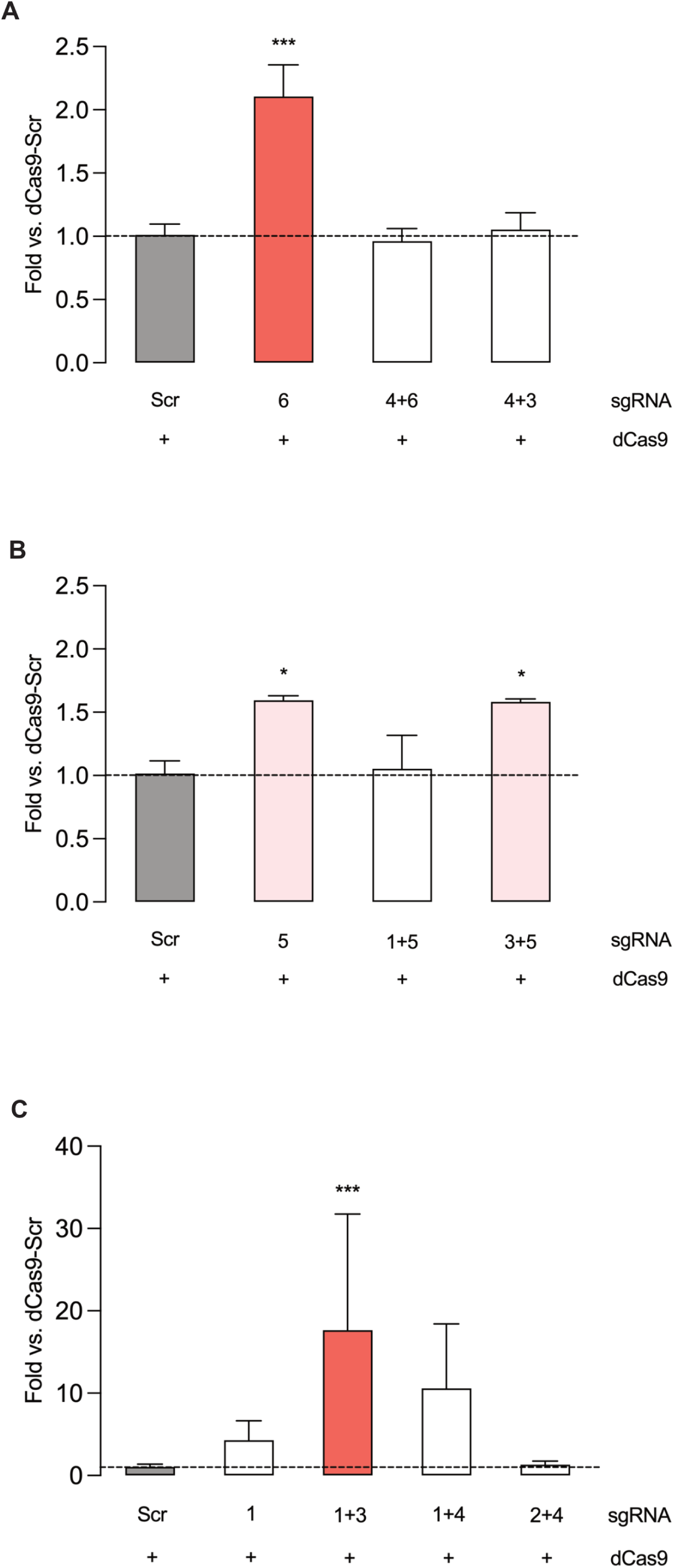
Activation of endogenous *Tfeb*, *Adam17*, and *Sirt1* expression. Gene expression analysis of *Tfeb* (A), *Adam17* (B), and *Sirt1* (C) in Neuro2A cultures transfected with respective sgRNA combinations along with MiniCas9V2. Samples were quantified using the expression of dCas9 as a baseline control. All values are presented as the mean ± SEM and quantified against a scrambled control (left, grey). For the histograms, bars with higher statistical significance are marked in darker shades of red. For multiple comparisons, *P≤0.05, **P≤0.01, ***P≤0.001, ****P≤0.0001.

In cells transfected with *Adam17* constructs, significant differences were observed between at least two groups when compared to the Scr baseline control (F (3,12) = 5.0, p<0.05) (Fig. 4B). Upregulation was observed in cells transfected with sgRNA 5 (1.60±0.03-fold) and sgRNA 3+5 (1.58±0.02-fold) respectively with expression 1.6-fold above baseline Scr control.

Finally, transfection with sgRNA 1+3, specific for *Sirt1,* revealed a statistically significant 17-fold increase in gene activation when compared to Scr baseline control (F (4,15) = 3.80), p<0.05) (17.67±7.04-fold) (Fig. 4C). In addition, a 10-fold increase in gene activity was detected in cells transfected with sgRNA 1+4, albeit the difference was not statistically significant.

### Tfeb activation using CRISPRa results in increased Lc3b levels

As a proof-of-principle final validation, LV expressing CRISPRa components were used to transduce NIH-3T3 cells. Subsequently, high content screening (HCS) was used to validate endogenous mouse *Tfeb* activation by directly quantifying Tfeb levels through immunofluorescence of individual transduced cells.

These experimental conditions resulted in a mean transduction efficiency of 72.8% across all wells. One-way ANOVA revealed significant differences in whole-cell Tfeb protein levels between groups (F (8,23) =30.49, p<0.0001) (Fig. 5A, C). Dunnett post hoc analysis against the Scr + MiniCas9V2 group revealed a slight Tfeb decrease in the groups expressing mRFP alone (0.86±0.01 fold). However, the groups receiving trehalose (1.25±0.02 fold), sgRNA 6 (1.24±0.04 fold) and sgRNA 4+6 (1.13±0.03 fold) exhibited significantly increased Tfeb expression.

**Figure 5:**
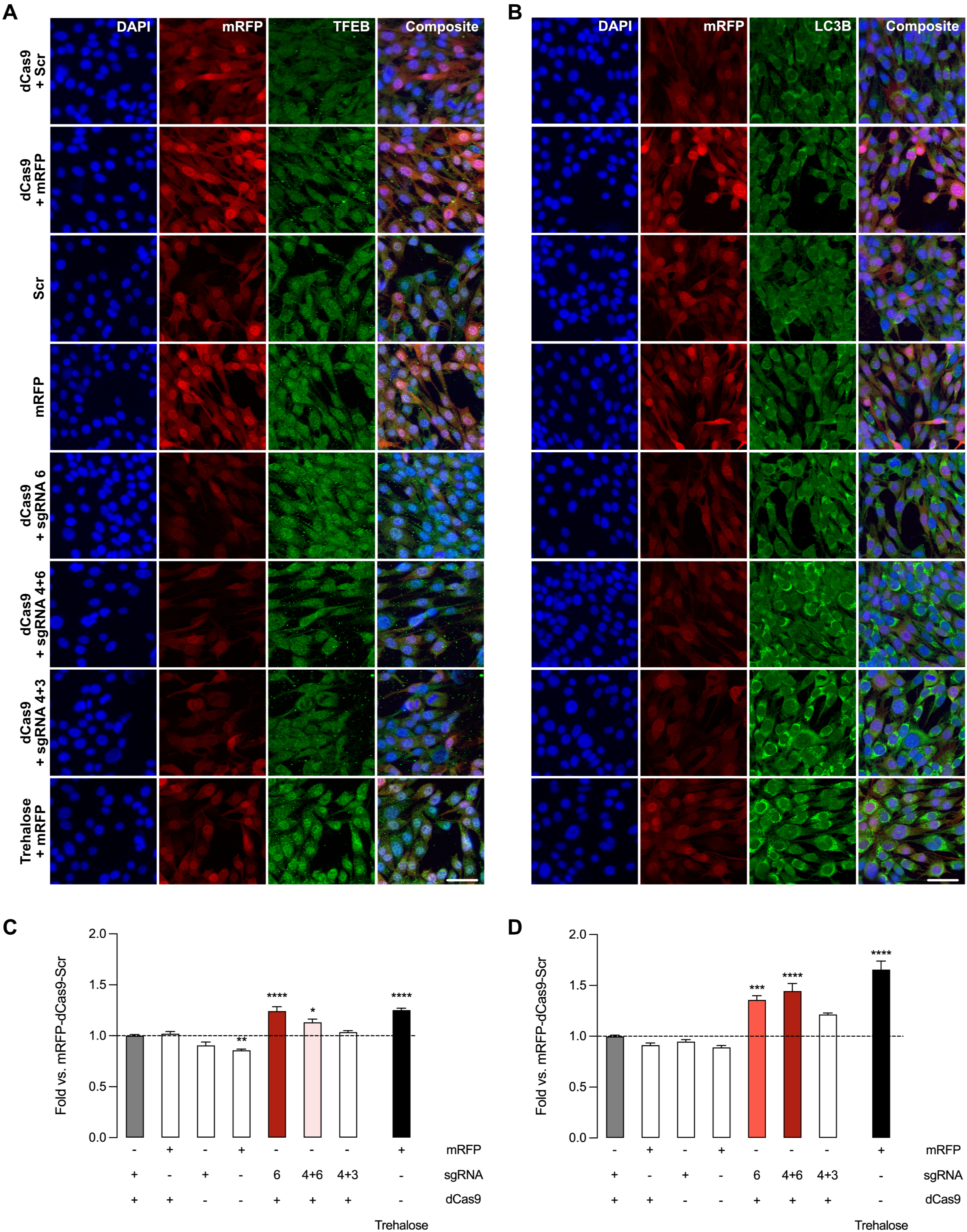
Transduction of NIH-3T3 cells with top sgRNA for *Tfeb* results in endogenous activation of *Tfeb* and increased Lc3b levels. NIH-3T3 cells were transduced with dual LVs at MOI 15. One LV expressed sgRNA and mRFP, whereas the second LV expressed MiniCas9V2. Fourty-eight hours after transduction, control cells were treated with trehalose. Seventy-two hours after transduction, cells were processed for immunofluorescence. A) Immunofluorescence against DAPI, mRFP fluorescence and *Tfeb* (Green). B) Immunofluorescence against DAPI, mRFP fluorescence and Lc3b (Green). C) Quantification of *Tfeb* protein levels. D) Quantification of Lc3b protein levels. Scale bar = 50 um. All values are presented as the mean ± SEM and quantified against a dCas9-Scr control (left, grey). For the histograms, bars with higher statistical significance are marked in darker shades of red. For multiple comparisons, *P≤0.05, **P≤0.01, ***P≤0.001, ****P≤0.0001.

Furthermore, autophagosomal marker Lc3b levels were quantified to assess if Tfeb activation increased lysosomal biogenesis (Fig. 5B, D). One-way ANOVA indicated significant group differences (F (8,23) = 39.0, p<0.0001). Dunnett post hoc analysis using the Scr + MiniCas9V2 group as control showed significant Lc3b increases in the groups receiving trehalose (1.66±0.08-fold), sgRNA 6 (1.36±0.04-fold) and sgRNA 4+6 (1.45±0.07-fold).

High-throughput screening enabled the assessment of whole cell Tfeb levels, indicating that sgRNA 6 and sgRNA 4+6 significantly increased endogenous Tfeb to levels comparable to those of the trehalose positive control. Concomitantly, Lc3b levels were also specifically increased in the sgRNA 6 and sgRNA 4+6 groups, once again to levels comparable with the trehalose positive control.

## 4 Discussion

This study describes the design and validation of an experimental screening assay to systematically assess sgRNA potential for CRISPRa applications using viral vectors. Furthermore, a cloning workflow was created to further facilitate the omnibus cloning of sgRNA into LV and AAV. The screening assay was validated with a miniature dCas9 coupled to a VPR system (Maria et al., 2020) to systematically screen sgRNA for three genes: *Tfeb, Sirt1* and *Adam17.* Activation of the promoter region proximal to the TSS was achieved in all genes tested, either by single sgRNA or dual sgRNA combinations. When transfected in Neuro2A cells, top single sgRNA or dual sgRNA combinations were able to activate endogenous *Tfeb, Sirt1* and *Adam17* transcription. As proof-of-principle, when delivered via LV, the validated sgRNA together with MiniCas9V2 were able to increase endogenous Tfeb levels, leading a concomitant increase in autophagosomes as measured by Lc3b levels. Thus, the screening assay was able to experimentally identify the minimal number of sgRNA needed for functional target gene activation.

Golden Gate (Engler et al., 2008) is a versatile cloning method that enables scarless, multitiered omnibus assembly of up to 35 DNA fragments (Bird et al., 2022). It is commonly used in CRISPR studies to insert 20-21 bp double stranded DNA, encompassing the DNA binding region, into sgRNA expression cassettes already present in final expression constructs (Cong et al., 2013; Savell et al., 2019). This simple design, while optimal for single one-step sgRNA cloning, lacks the flexibility to switch or rearrange multiple sgRNA cassettes if needed. For this reason, Kabadi and colleagues developed a two-tier Golden Gate system where double-stranded DNA is first cloned and inserted into a plasmid containing one sgRNA expression cassette (Kabadi et al., 2014). A second Golden Gate reaction is then used to insert up to four different sgRNA cassettes into lentiviral transfer vectors containing an active Cas9 for gene editing. As this Golden Gate design uses different pol III promoters for each of the sgRNA cassettes, each short RNA will be expressed with different efficiencies (Goguen et al., 2021), potentially causing suboptimal activation, especially for multiplex activation. More recently, Savell and colleagues used Golden Gate to clone up to eight sgRNA cassettes in tandem, all driven by separate human U6 promoters, for multiplex CRISPRa using a dual lentiviral vector system (Savell et al., 2020). For our research needs, we required a flexible cloning workflow where sgRNAs would be first validated in a screening assay and subsequently cloned and inserted into an LV or AAV CRISPRa system. Although L3G0 cloning can assemble full sgRNA cassettes similarly to the systems described above, it differs in two key points. First, in the L3G0 workflow, the sgRNA cassettes are first screened before being cloned into viral vectors, in contrast with Savell et al, where the sgRNA are first cloned into an LV for individual screening and subsequently multiple sgRNA are re-cloned into a second LV for multiplex CRISPRa. Second, the L3G0 workflow combines up to eight sgRNA cassettes using multiple Golden Gate destination sites with the flexibility of subcloning these cassettes into LV and AAV. This workflow is designed to minimize the time and steps needed from initial sgRNA screening to final incorporation of validated sgRNA into viral vectors for in vivo or ex vivo gene therapy applications.

The design of effective sgRNAs is a key factor for successful CRISPRa applications (Kampmann, 2018; Casas-Mollano et al., 2020; Pandelakis et al., 2020; Liu et al., 2021). Commonly used tools such as CHOPCHOP (Labun et al., 2019) can be complemented further by machine learning to predict sgRNA function for CRISPRa applications (Horlbeck et al., 2016). However, designing and selecting sgRNA for CRISPRa using only bioinformatic tools has drawbacks. First, different Cas-proteins bind optimally at different promoter regions. For example, Cpf1 can activate genes by binding −600 to +400 from the TSS, in contrast with SpCas9, which uses +400 to 0 from the TSS (Gilbert et al., 2014; Tak et al., 2017; Chen et al., 2020). In addition, these predictive tools have been developed for SpCas9 (Horlbeck et al., 2016) and it is currently unclear how suitable these tools are for other Cas proteins. Hence, most CRISPRa studies either employ multiple sgRNA together without prior validation of activation potential (Kunii et al., 2018; Weltner et al., 2018; Choi et al., 2020) or select few sgRNA and assess their effectiveness experimentally (Cheng et al., 2013; Maeder et al., 2013; Savell et al., 2020; Schoger et al., 2020).

Using large cassettes containing multiple sgRNAs without any prior selection is suboptimal for gene therapy for several reasons. First, the inherent size constrains for cargo in viral vectors makes cassettes with excessive numbers of sgRNAs incompatible with viral vector delivery. Moreover, using multiple sgRNA increases the likelihood of off-target effects, and may also increase the chance of recombination events and deficient viral production (Casas-Mollano et al., 2020; Schoger et al., 2020). Importantly, multiple sgRNA usage does not guarantee robust gene activation and may result in reduced target gene expression, despite being common practice in CRISPRa applications (Cheng et al., 2013; Schoger et al., 2020).

Data from the experimental screening assay indicated that using a single sgRNA does not necessarily lead to detectable promoter activation. This is in line with data from multiple in vitro and in vivo studies where not all genes were able to be activated with one sgRNA (Weltner et al., 2018; Savell et al., 2020; Schoger et al., 2020). In addition, our data show that similar to single sgRNAs, not all dual sgRNA combinations lead to activation. Again, this is in line with other studies that observed a lack of gene activation with several dual or multiple sgRNA combinations (Lin et al., 2015; Schoger et al., 2020). The maximum level of activation observed in the screening assay was also dependent on the promoter targeted, with mouse *Tfeb* and *Sirt1* resulting in 4-to-5-fold activation, whereas mouse *Adam17* led to 7-fold activation of their respective promoters. Two important conclusions can be drawn from this set of data. First, not all individual sgRNA will cause gene activation. Second, given the preferentially antagonistic interaction between sgRNA, arbitrary selection without validation will likely result in false negatives. These points further highlight the importance of assessing such interactions before experimental applications. Taken together, the data from our study and previous literature suggest that the optimal number of sgRNAs required for robust CRISPRa activation is gene specific. Therefore, selecting and experimentally screening for the minimum amount of sgRNA/cRNA for CRISPRa is needed for optimal results.

When examining the effective relationships between sgRNA candidates, antagonistic interactions were observed more frequently than synergistic interactions. However, it remains important to assess the interactions between the respective sgRNA candidates along with the subsequent level of activation. As an example, in the screening assay for *Adam17,* sgRNA 4 displayed antagonism when coupled with either sgRNA 1 or sgRNA 5. When comparing sgRNA 1+4 to sgRNA 1+5 (Fig. 3B), sgRNA 1+4 displays a complete lack of activation along with a strong antagonistic interaction. Looking further, sgRNA 1 alone has a statistically significant activation which appears nullified when combined with sgRNA 4. This suggests that the antagonistic response of sgRNA 4 is enough to abolish the otherwise effective candidate sgRNA 1. However, in sgRNA 1+5, the detected antagonism coupled with a strong activation suggests that the gene may already operate at a maximum level of activity due to sgRNA 5, and is not nullified by the seemingly antagonistic interaction with sgRNA 1. These observations underline the importance of considering sgRNA antagonism in concert with the resulting gene activity response.

Studies validating sgRNA often use qPCR to test between two and seven sgRNAs (Konermann et al., 2015; Kunii et al., 2018; Colasante et al., 2019, 2020; Savell et al., 2019, 2020; Choi et al., 2020). In contrast to these studies, we opted to screen sgRNA using a fluorescence-based assay. In addition to allowing a readout at protein level by quantifying transgene expression based on TdTomato, fluorescence-based methods are cost-effective and allow easy scaling-up of sgRNA screening when compared to qPCR methods. We started by developing fluorescence-based tools for the assessment of up to four sgRNAs (Maria et al., 2020). Moving forward, we further optimized the assay conditions and integrated the assay into a cloning workflow, allowing us to screen up to 28 separate single- or combined sgRNAs in a 96-well plate format. This let us systematically assess sgRNA efficacy throughout promoter regions relevant to MiniCas9V2.

From inception, the screening assay was designed to test sgRNAs/cRNAs for any gene or Cas protein. However, the current iteration of the screening assay has limitations. As the assay is plasmid-based, it does not consider gene and cell-specific epigenetics. For this reason, top-selected sgRNAs may need to be further tested on endogenous genes in order to fully determine their potential for gene activation. Nevertheless, it remains unclear what impact epigenetic modifications have on CRISPRa as effective sgRNAs can activate the same genes across different cell types, whether in culture or in vivo (Cheng et al., 2013; Maeder et al., 2013; Chavez et al., 2015; Kiani et al., 2015; Vora et al., 2018; Zhou et al., 2018; Colasante et al., 2019, 2020; Matharu et al., 2019; Savell et al., 2019, 2020; Maria et al., 2020; Schoger et al., 2020; Giehrl-Schwab et al., 2022).

In the *Adam17*-screening assay, we observe similarly significant levels of gene activation between sgRNA 5, sgRNA 1+5, and sgRNA 3+5 respectively. Interestingly, at mRNA level, we observe a significant increase in *Adam17* expression in cells transfected with sgRNA 5 and sgRNA 3+5 respectively, but not with sgRNA 1+5. A similar trend was observed in the *Tfeb* assay where sgRNA 6, sgRNA 4+6, and sgRNA 4+3 all displayed significant upregulation, but solely sgRNA 6 resulted in a significant activation at mRNA level. This may suggest a discrepancy in sgRNA efficacy when targeting a fully accessible promoter, such as from a transfected plasmid, as opposed to an endogenous target. Additionally, sgRNAs may act individually to a greater extent in reporter models with a high number of available transgene copies, as opposed to in endogenous genes where only two target copies are available for regulation, therefore promoting a different kinetic model (Duarte et al., 2023). However, the increase in mRNA levels observed in this study were comparable to studies targeting therapeutic genes for CRISPRa (Chavez et al., 2015; Colasante et al., 2019, 2020; Savell et al., 2020). It is therefore important to consider that upregulation of therapeutic targets generally entails activating genes that have minimal constitutive expression. Thus, the activation levels, especially at protein level, may not be dramatically increased, unlike proof-of-concept genes that may be more easily targeted and regulated.

Given the variation in expression ceilings for different genes, a functional readout may be required in addition to bulk expression analysis. *Tfeb* is one such example, as it is a highly regulated transcription factor with therapeutic potential in several neurodegenerative diseases (Martini-Stoica et al., 2016; Chen et al., 2021). When transfected into Neuro2A cultures (Fig. 4), a significant increase in *Tfeb* activity was observed only when sgRNA 6 alone was applied. However, transduction of MiniCas9V2 together with sgRNAs 6 or 4+6 in culture resulted in increased *Tfeb* levels and was sufficient to induce autophagosome formation, indicating that activation of *Tfeb* was functional (Fig. 5). This points to a divergence in assessing functional response between transfection- and transduction-based methods and may prove important when determining the most reliable approach to assess the functional readout for a gene of interest (Duarte et al., 2023). In this case, the rapid transcriptional response associated with transfection-based methods may not be desirable when assessing the effects from upregulation of the *Tfeb* gene.

Interestingly, while sgRNA 6 alone indicated a higher recorded level of *Tfeb* activation than sgRNA 4+6, the latter appeared to result in a higher expression of the downstream effector Lc3b. Considering the nature of *Tfeb* as a highly regulated transcription factor with broad effects on cellular homeostasis (Napolitano and Ballabio, 2016) (Chen et al., 2021), the highest level of activation possible may not be desirable for an optimal therapeutic response. Additionally, given the tight regulation of *Tfeb* under healthy cellular conditions, the functional readout of *Tfeb* activation may be more efficiently studied in disease models of protein aggregation to determine potential therapeutic effects. Therefore, we aim to investigate the effects of *Tfeb* activation using CRISPRa in further studies.

In summary, we describe a systematic experimental approach to screen sgRNA/cRNA for CRISPRa studies. Our assay is designed to be universal and has allowed the identification of efficient sgRNAs for all genes tested in this study. The multiplexing potential of this workflow may assist in accelerating and streamlining future developments of CRISPRa-focused applications.

## 6 Author Contributions

EA, DL, ES, FD, RN and LQ contributed to the experimental design, performed the experiments, and analysed the data. EA, DL, ES and LQ produced the lentiviral vectors. FD and LQ contributed to establishing the initial sgRNA screening assay. LQ oversaw, designed, and led the experiments. EA and LQ wrote the manuscript. CL secured funding, oversaw and coordinated the project. All authors contributed to manuscript revision, read, and approved the submitted version.

## 8 Acknowledgments

We are grateful to Marie Persson Vejgården for laboratory assistance, and to Anna Hammarberg and the Multipark infrastructure at Lund University for availability and assistance with the Operetta CLS.

## Supplementary Material

**Supplementary Figure 1:**
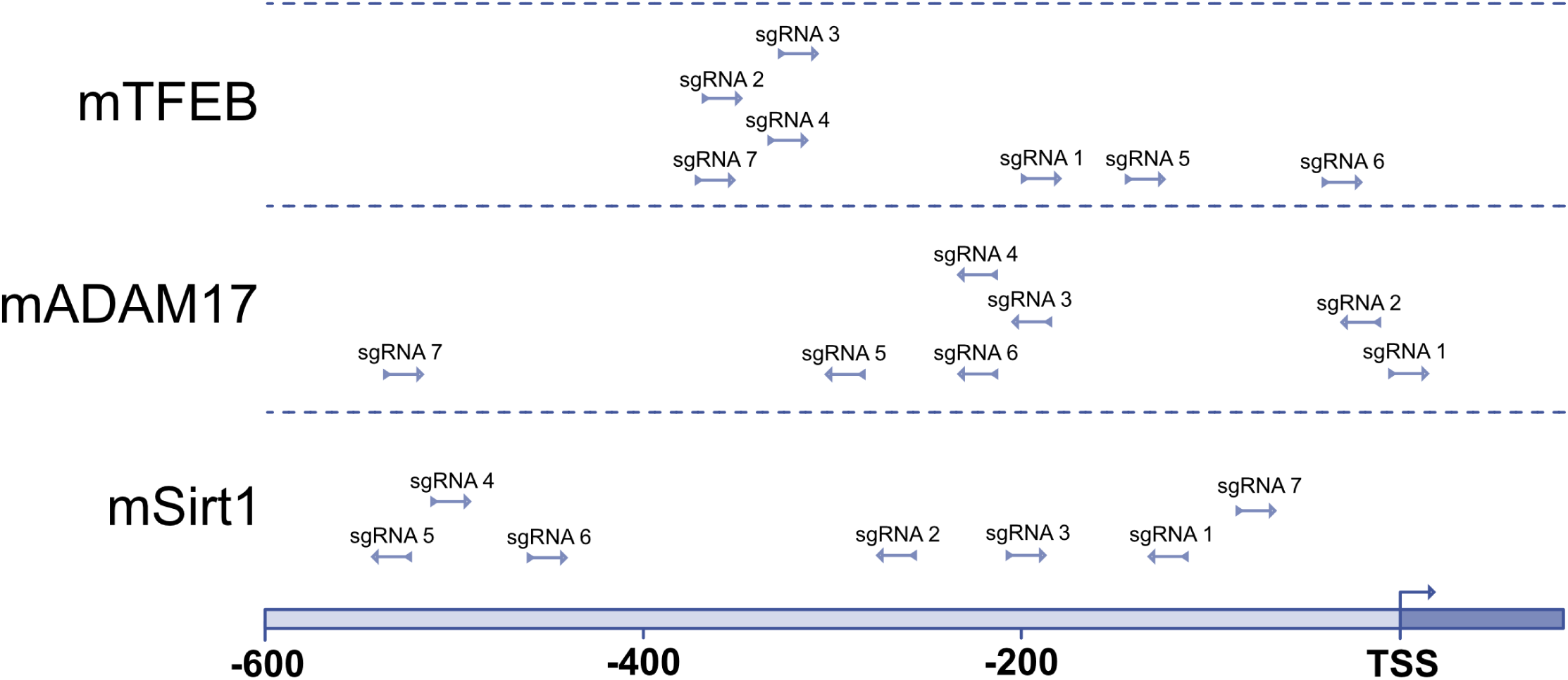
sgRNA sequences relative to transcriptional start site. sgRNA 1 to 7 for mTFEB, mADAM17 and mSIRT1 placed relative to the transcriptional start site (TSS) for each of their respective genes.

**Supplementary Figure 2:**
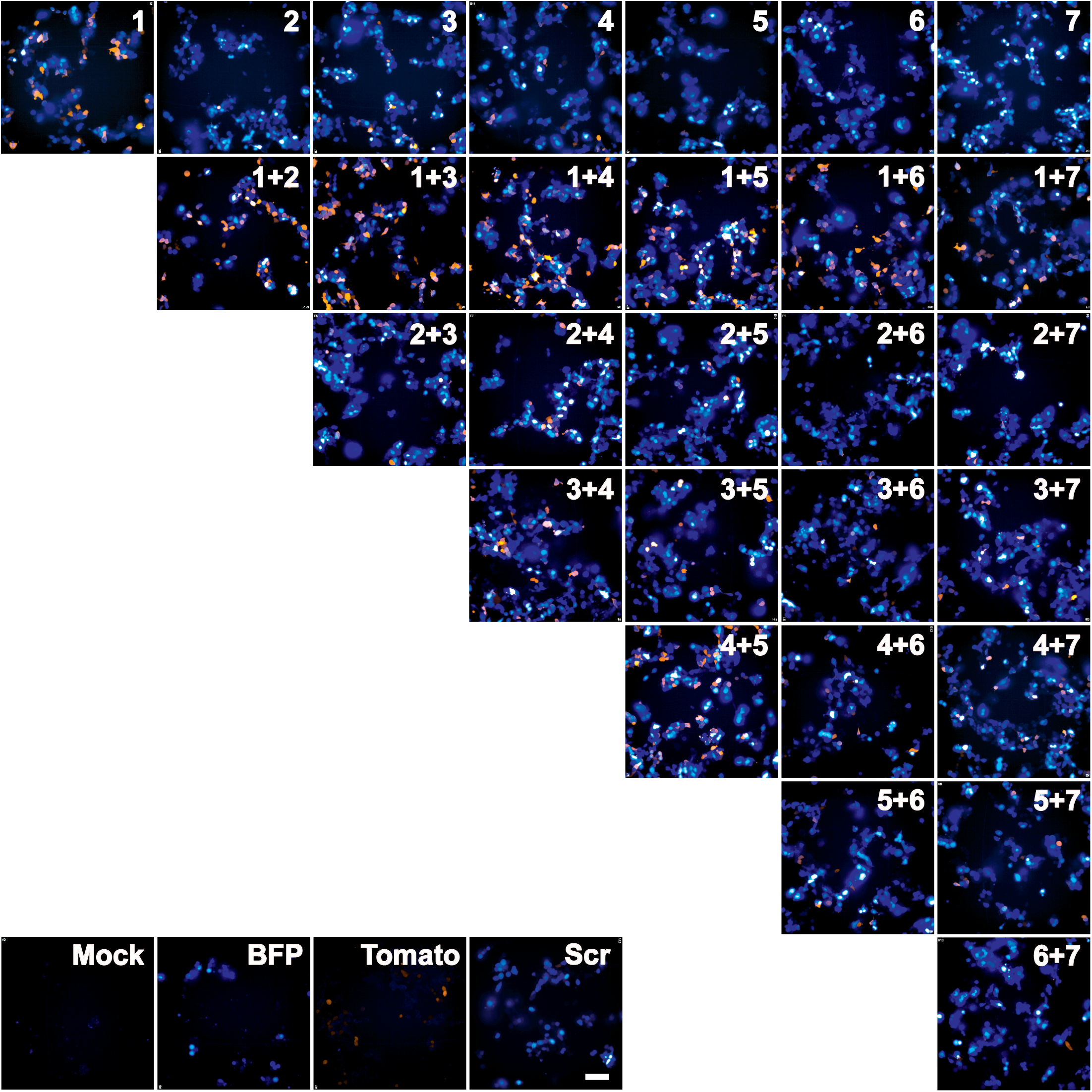
Fluorescence images of the screening assay using mSIRT1 as an example. 293T cells were transfected with pDPL0 expressing sgRNA, reporter constructs expressing TdTomato, MiniCas9V2 and BFP. Forty-eight hours after transfection, the cells were fixed and the TdTomato and BF fluorescence were measured. Each number denotes an sgRNA. Mock-mock transfected cells. BPF-Cells transfected with BFP and MiniCas9V2. Tomato-Cells transfected with TdTomato reporter and MiniCas9V2. Scr-Cells transfected with Scramble sgRNA, MiniCas9V2, TdTomato reporter and BFP. Scale bar-100 μm.

**Supplementary Table 1:** Settings for immunofluorescence cell detection in Operetta CLS.

|  |  |  |
| --- | --- | --- |
| 1 | FIND NUCLEI | • DAPI channel<br>• Method B<br>• >70um<sup>2</sup><br>• Individual threshold = 0.45<br>• Contract >0.1 |
| 2 | FIND CYTOPLASM | • mRFP channel<br>• Nuclei<br>• Method B |
| 3 | CALCULATE MORPHOLOGY PROPERTIES | • Area<br>• Roundness<br>• Ratio: Width/Length |
| 4 | SELECT POPULATION | • Nucleus<br>• Area >400 um<sup>2</sup><br>• Roundness: 0.5<br>• Width/length ratio: 0.45 |
| 5 | SELECT POPULATION (2) | Nuclei selected → Common filters → Remove border objects |
| 6 | CALCULATE INTENSITY PROPERTIES | mRFP (based on (5)) |
| 7 | CALCULATE INTENSITY PROPERTIES (2) | Alexa 647 in cell (based on (5)) |
| 8 | CALCULATE INTENSITY PROPERTIES (3) | Alexa 647 in nucleus (based on (5)) |
| 9 | CALCULATE INTENSITY PROPERTIES (4) | Alexa 647 in cytoplasm (based on (5)) |
| 10 | SELECT POPULATION | (5) → filter by property<br>→ intensity of nuclear mRFP, mean >600<br>(determined cut-off through scatterplot) |

